# Population density is a selective driver of urban soil microbiomes: evidence from bacteria, fungi and protozoan communities in a regional city

**DOI:** 10.64898/2026.09.20.752949

**Authors:** Jessica Grierson, Penelope Jones, Andrew Bissett, Shane Powell, Emily J Flies

**Author notes:** Corresponding author: Jessica Grierson, University of Tasmania, Private Bag 55, Hobart, Tasmania 7001.

## Abstract

Urbanisation is widely recognised as a major driver of environmental change, yet its impacts on soil microbiomes remain poorly understood, particularly outside megacities and beyond bacterial taxa. Population density is frequently used as a proxy for urban intensity, but its relevance across different microbial kingdoms and urban contexts is unclear. Here, we investigated how soil microbial communities vary across population density bands in the regional, temperate city of Hobart, Australia. Using a multi-kingdom metabarcoding approach, we analysed bacterial, fungal, and protozoan communities from soils collected in urban green spaces along a population density gradient.

Bacterial alpha-diversity differed significantly across population density bands, exhibiting an inverse U-shaped relationship, with the highest diversity at green spaces in intermediate population density areas. In contrast, fungal and protozoan alpha-diversity did not vary with population density. Community composition (beta-diversity) did not differ consistently across density bands for any microbial group, and variation in soil physicochemical properties explained more community variation than population density alone. These results indicate that population density is a selective, rather than universal, driver of urban soil microbiomes and that responses to urbanisation differ markedly among microbial kingdoms.

Our findings highlight important limitations of using population density as a sole indicator of urban impact and challenge the generalisability of conclusions drawn from bacteria-focused studies in large cities. By demonstrating kingdom-specific and context-dependent responses in a regional city, this study underscores the need for multi-kingdom, multi-scale approaches to better understand how urbanisation shapes soil microbiomes and their potential implications for urban ecosystem functioning and human health.

## 1 Introduction

The world has become increasingly urban over recent decades, with the proportion of the global population living in cities more than doubling from 20% in 1950 to 45% in 2025 (United Nations, 2025). This continuing urbanisation creates a clear imperative to understand the implications for human health. Notably, some non-communicable diseases, including allergies, asthma, as well as other immune-mediated conditions, are more prevalent in urban areas, leading to concerns that their burden may increase as urbanisation progresses (Flies et al., 2019). The links between non-human nature and human health – including in the context of non-communicable diseases - are numerous (Hartig et al., 2014) but one proposed thread is exposure to diverse microbiomes (Flies et al., 2019; Flies et al., 2018).

Urban soils have important impacts on human health through multiple pathways, including by providing a source of exposure to biodiverse environmental microbial communities (Sun et al., 2023). Public and residential green spaces are key spaces for urban populations to encounter biodiverse microbial communities from soil, air and other surfaces (Matthews et al., 2024). However, urban soil microbiomes experience homogenisation at the global scale, where the same suites of taxa/traits recur across cities (Delgado-Baquerizo et al., 2021; Zhang et al., 2025). Therefore, there is a need to understand the extent to which urban green spaces can offer exposure to microbial diversity, and the factors which impact urban soil communities.

To date however, outdoor urban microbiomes (e.g. soil, air, water, surfaces) remain comparatively understudied and less comprehensively characterised relative to other microbiome communities (Masson et al., 2025; Sessitsch et al., 2023). Although urbanisation has been shown to influence environmental microbiomes (Flies et al., 2020; Wang et al., 2018; Yan et al., 2020) the direction and magnitude of these effects remain highly context dependent. Urban soil microbial communities are shaped by multiple interacting environmental and anthropogenic factors, including soil physicochemical properties, vegetation characteristics, land-use history and management practices (Islam et al., 2020; Thompson et al., 2023; Zhang et al., 2019). Consequently, urban soils are increasingly recognised as heterogeneous ecosystems rather than uniformly degraded systems.

Many existing studies compare soil microbial communities between urban and rural environments (Abrego et al., 2020; Yan et al., 2020) or across urbanisation gradients (Gao et al., 2023; Li et al., 2023; Qu et al., 2025; Rusterholz & Baur, 2023). These approaches have identified important roles for edaphic variables such as soil pH, nutrient availability, organic carbon and moisture, together with built-environment characteristics including impervious surface cover, in explaining microbial variation. However, comparisons across contrasting landscapes or regions are often confounded by differences in climate, vegetation, land-use history and management, making it difficult to disentangle the specific effects of urbanisation (Nugent & Allison, 2022). Without pre-urbanisation baseline data, understanding the effects of urbanisation on soil microbiomes requires spatial comparisons using proxies of urbanisation intensity (Masson et al., 2025; Nugent & Allison, 2022).

Population density is frequently used as a proxy for urban intensity because it can capture aspects of human pressure associated with infrastructure density, soil disturbance, and management intensity. Studies using population density or related urban intensity metrics have demonstrated that increasing human density can influence soil microbial communities through changes in soil properties, resource availability, disturbance regimes and habitat conditions (Liddicoat et al., 2016; Mills et al., 2019; Wang et al., 2018). However, responses of bacterial diversity to population density have been inconsistent, with studies reporting increases, decreases or no significant changes in diversity across urban gradients (Flies et al., 2020; Liddicoat et al., 2016; Mills et al., 2019; Wang et al., 2018). Our previous work demonstrated that bacterial diversity within regional city parks varied with intensity of human use (Grierson et al., 2023), highlighting the importance of local environmental context. Together, these findings suggest that the relationship between population density and soil microbiomes is highly context dependent, and that the mechanisms linking human density to microbial diversity remain unresolved.

In addition, most urban green space microbiome studies focus primarily on bacterial communities (Masson et al., 2025; Tan et al., 2019; Wang et al., 2018; Yan et al., 2016; Yan et al., 2020; Zhang et al., 2019). However, our earlier work has shown that bacteria, fungi and protozoans in soil respond differently to urban living (Grierson et al., 2023). Thus, a major literature gap exists with respect to the impact of urban-associated environmental factors on other microbiome components such as fungi, and protozoa (Sun et al., 2023). Consequently, multi-kingdom studies are required to determine how different components of the soil microbiome respond to urbanisation.

Much of the existing urban soil microbiome literature has been conducted in large metropolitan or megacity contexts, where urbanisation is often intense and associated with extensive landscape modification (Masson et al., 2025; Nugent & Allison, 2022). However, a substantial proportion of the global urban population resides in small and medium-sized cities (United Nations, 2025). Urbanisation within smaller and regional cities can follow different spatial and developmental trajectories from those of highly urbanised metropolitan regions, with differences in rates of land expansion, population density and landscape transformation (Boone et al., 2012). These differences raise the possibility that environmental pressures and ecological drivers identified in highly urbanised environments may not yet operate, or may operate differently, areas of less intense urban development. Yet relatively little is known about how soil microbiomes respond across different levels of urban development (Masson et al., 2025; Nugent & Allison, 2022). Understanding soil microbiomes in regional cities is therefore critical for developing broadly applicable approaches to urban planning, biodiversity conservation, and ecosystem management.

This study seeks to address these gaps with a multi-kingdom study of the soil microbiome along urban density bands in Hobart, Australia. Hobart is an Australian regional capital city (city-centre population 55 977; Greater Hobart area population 254 930) (Australian Bureau of Statistics, 2025; City of Hobart, 2025) on the island of Tasmania and represents an important opportunity to consider the impacts of urbanisation in a relatively small, low population density city. Our study focuses on the soil microbiomes of two urban green spaces where people interact with soils: public playgrounds and private backyards and considers the composition and diversity of three major microbiome taxa: bacteria, fungi, and protozoa. Parks and backyards are two common green spaces used by urban residents, and we have found that these two spaces have similar diversity and composition (Grierson et al., 2023). We now explore whether communities in these green spaces are different across population density bands.

This study investigates whether population density is associated with soil microbial diversity and community composition across bacteria, fungi, and protozoa within a regional city. By adopting a multi-kingdom approach in a small urban centre, this study addresses three major gaps in urban microbiome research: over-reliance on bacterial communities, the limited understanding of population density as an urbanisation metric for microbial communities, and the scarcity of studies from regional cities.

## 2 Methods

### 2.1 Study region

This study builds on the work published under Grierson et al. (2023), using the same soil samples to further explore soil microbial communities across the greater Hobart region. These soil samples were collected across Hobart (the Greater Capital City Statistical Area; 42.88° S, 147.33° E) between the 4^th^ and 22^nd^ of November 2019 (Figure 1). Hobart has a temperate maritime climate, with mean daily temperatures ranging from 7 to 17 °C across the year and mean annual rainfall of 612 mm (Bureau of Meteorology, 2022). Hobart lies under kunanyi/Mount Wellington and along the Derwent River, and is built upon multiple soil types, including dolerite, mudstone, sandstone, and alluvial soils. The city mainly comprises suburbs with relatively low population density (182–1258 people/km^2^), with the inner city achieving maximum population densities of 1694/km2 (average density 1518 people/km^2^) (see Figure 1).

**Figure 1.**
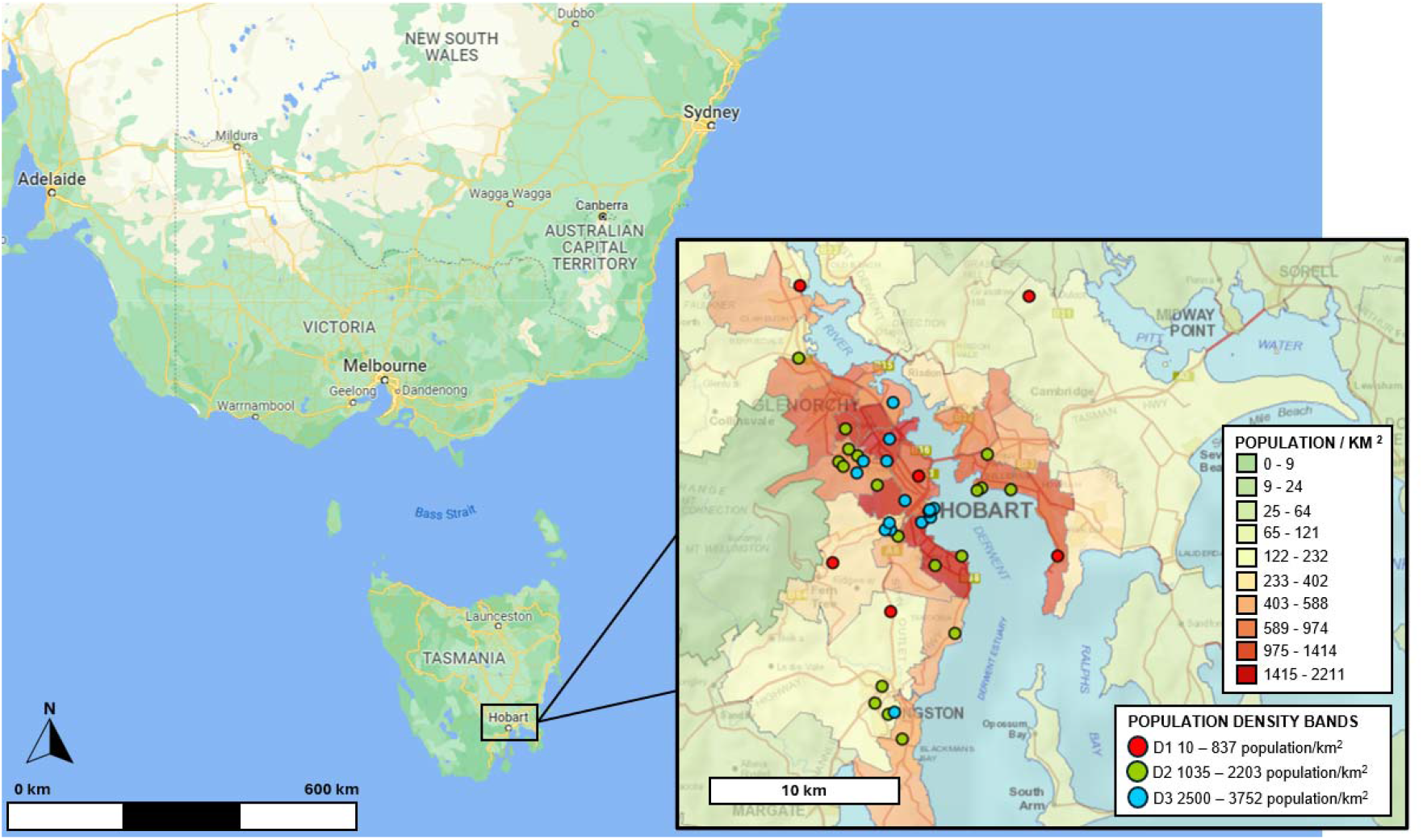
Map of population density band sampling locations layered across a population density map of Hobart, Australia. Shown are population density bands: D1 = 10 – 837 population/km^2^ (red), D2 = 1035 – 2203 population/km^2^ (green), and D3 = 2500 – 3752 population/km^2^ (blue). Background shading shows population density (people/km^2^) across the greater Hobart region. Information was sourced from ListMAP (Land Tasmania, 2022), according to the Australian Bureau of Statistics 2016 census data (Australian Bureau of Statistics, 2017).

### 2.2 Soil sampling

Soil samples were collected along a population density gradient, and sites were stratified across clear break points in the population density gradient (Australian Bureau of Statistics, 2017) to create three population density bands: D1 (10 – 837 population/km^2^), D2 (1035 – 2203 population/km^2^) and D3 (2500 – 3752 population/km^2^; Figure 1). Soil samples were collected from both private backyards (N=29) and public parks with playgrounds (N=12; in areas of high intensity of human use e.g., around seating (benches, picnic tables), walkways (between pathways, gates) and around playground areas). Our previous work found that soil microbial communities across these two urban green spaces were not significantly different (Grierson et al., 2023). Thus, this work combined these datasets to explore soil microbial communities across population density bands. The private backyard sampling sites were dependent on volunteer recruitment, so the distribution across population density bands and soil types (Islam et al., 2020) was not even (Figure 1, Figure S1, Table S1). Permission from the relevant council was received prior to soil collection for all public park sampling; participant consent was received prior to soil collection from backyards, and for the use of their information (Ethics ref. H0018356).

### 2.2 Soil collection and physicochemical analysis

Soil samples and site-specific metadata were collected according to the Australian Microbiome (AM) protocols (Mazard et al., 2019). Soil samples were collected across a 25 x 25m area of lawn, isolated from other vegetation to minimise the impact of vegetation differences on microbial compositions (Islam et al., 2020). Nine to ten sub-samples were collected across the entire grid from the soil surface (A) horizon (0-10cm depth) and combined to create a pooled sample for each site. The pooled sample was stored at -80°C and sent to the Australian Genome Research Facility (ARGF) laboratory (Adelaide, SA) for DNA extraction and sequencing analysis. The remaining sample was air dried and sent to the CSBP Soil and Plant Laboratory (Perth, WA) for physicochemical analysis according to AM protocol (CSBP, 2019; Mazard et al., 2019). The edaphic properties measured are as listed on Table S2. Results of physicochemical analysis found that one ammonium nitrogen and four nitrate nitrogen samples were below the limit of detection (1 mg/kg) and were assigned a value of 0.5 mg/kg (range of ammonium nitrogen 1-148 mg/kg and nitrate nitrogen 1-67 mg/kg).

### 2.3 DNA extraction, amplification, and sequencing

Soil DNA extraction, amplification, and sequencing were conducted at the ARGF laboratory, following the AM protocol (Mazard et al., 2019) and full descriptions of the methods are described in Grierson et al. (2023). In short, DNA extraction was completed in triplicate (0.25g of soil used) and combined to give a single DNA sample used for further analysis. Soil bacteria, fungi, and microbial eukaryotes were identified by meta-barcoding through paired-end sequencing on the Illumina MiSeq® platform (Illumina Inc.; Mazard et al., 2019). Bacterial communities were identified through amplification of the 16S rRNA gene (V1-V3) using primers 27F (Lane, 1991) and 519R (Lane et al., 1985). Fungal communities were identified through amplification of the ITS gene using primers ITS1F (Gardes & Bruns, 1993) and ITS4 (Gardes & Bruns, 1993; White et al., 1990). Eukaryotes were identified by amplification of the 18S rRNA gene (V9) using primers 1391F and EukBR (Amaral-Zettler et al., 2009).

### 2.4 Sequence analysis and bioinformatics

Bioinformatic analysis was undertaken by AM, following their standard amplicon workflow (Mazard et al., 2019; Ostrowski, 2019). Amplicon paired ends were merged and denoised to zero radius Operational Taxonomic Units (zOTU’s) at 97% similarity and classified with taxonomy using the RDP Bayesian classifier (Wang et al., 2007). 16S rRNA and 18S rRNA sequences were matched to the Silva database (Chuvochina et al., 2026), and ITS sequences matched using the UNITE SH database (Nilsson et al., 2019), with a probability cut-off of 60%.

Only 16S rRNA sequences from bacteria were included in the bacterial dataset. Fungal ITS sequences “unclassified” at Phylum level were discarded. 18S rRNA sequences classified as animal (excluding nematodes), plant, fungal or “unclassified” at Phylum level were removed from all analyses, which involved the removal of 11,855 sequence reads (82 % of the total - 18S rRNA zOTU reads). zOTUs classified as “Helminths” were kept in the analysis within the “Protozoan” group.

### 2.5 Statistical analysis

All statistical analyses were conducted in R version 4.3.1 (R Core Team, 2020) using RStudio (RStudio Team, 2020).

#### 2.5.1 Alpha-diversity

The first determination of soil diversity across the population density bands was through the calculation of alpha-diversity using zOTU data. Prior to the calculation of alpha-diversity, zOTU count data were rarefied to sufficient sampling depth based on rarefaction curves (bacteria = 38 792 reads, fungi = 45 998 reads, eukaryotes = 1 536 reads; Table S2) using the *rarefy_even_depth* function in the *phyloseq* package (McMurdie & Homes, 2013).

Alpha-diversity was assessed by calculating (a) Total Species Richness, using the *specnumber* function in the *vegan* package, and (b) Shannon-Weaver Diversity index, using the *alpha* function in the *microbiome* package (Oksanen et al., 2019). Differences in alpha-diversity indices were assessed using one-way ANOVA with the factor of population density (three levels) and Tukey HSD multiple comparisons after checking for normality and homogeneity of variance (Table S3). A Spearman’s rank order correlation matrix was computed using *rcorr* function in the *Hmisc* package (FE Harrell Jr & Dupont, 2021) to observe any relationships between alpha-diversity indices and soil environmental variables across the population density bands.

#### 2.5.2 Beta-diversity

We explored beta-diversity using a compositional data analysis (CoDa) approach, using transformed zOTU data and computing Aitchison distance (Gloor et al., 2017; Quinn et al., 2019). In short, zero counts were replaced using the *cmultrepl* function in the *zCompositions* package (Palarea-Albaladejo & Martin-Fernandez, 2015), and the dataset was *centered log-ratio (clr)* transformed, using the *clr* function in the *compositions* package (Boogaart et al., 2020). Aitchison distance was determined by calculating Euclidean distance on the *clr* data, using the *distance* function in the *phyloseq* package (McMurdie & Holmes, 2013).

PERMANOVA was used to test differences between population density bands, using *adonis2* function in the *vegan* package (Oksanen et al., 2019), with equal dispersion among groups tested for using the *betadisper* function also in the *vegan* package (Oksanen et al., 2020). Principal component analysis, based on Aitchison distance, was used to visualise the dissimilarity between sites based on community composition. Environmental variables significant in explaining the variation of the ordination axes (Bonferroni-adjusted significance threshold < 0.05) were standardised using *decostand* function, and fitted onto the ordination using *envfit* function, in the *vegan* package (Oksanen et al., 2019).

#### 2.5.3 Community composition

We observed broad community composition using stacked bar charts at the phylum level across population density bands. For the dominant phylum groups, the bar chart was expanded to show the top families present within those phyla to aid visualisation. Families were plotted if they met the threshold of a relative abundance greater than: 5% for bacterial, 20% for fungal, and 10% for protozoan families.

## 3 Results

### 3.1 Alpha-diversity

There were significant differences in bacterial alpha-diversity only, across the population density bands (ANOVA, Total Species Richness, p = 0.004, Shannon-Weaver Diversity, p = 0.014, Figure 2, Table S4). Alpha-diversity of soil bacteria followed an inverse U-shaped trend between groups, where diversity was significantly higher in the middle population density band (D2) compared to low population density (D1; Tukey HSD, Total Species Richness, p = 0.039) and higher population density (D3; Tukey HSD, Total Species Richness, p = 0.008, Shannon-Weaver Diversity, p = 0.017). There were no significant differences in alpha-diversity between population density bands for soil fungi and protozoa (Figure 2, Table S4).

**Figure 2.**
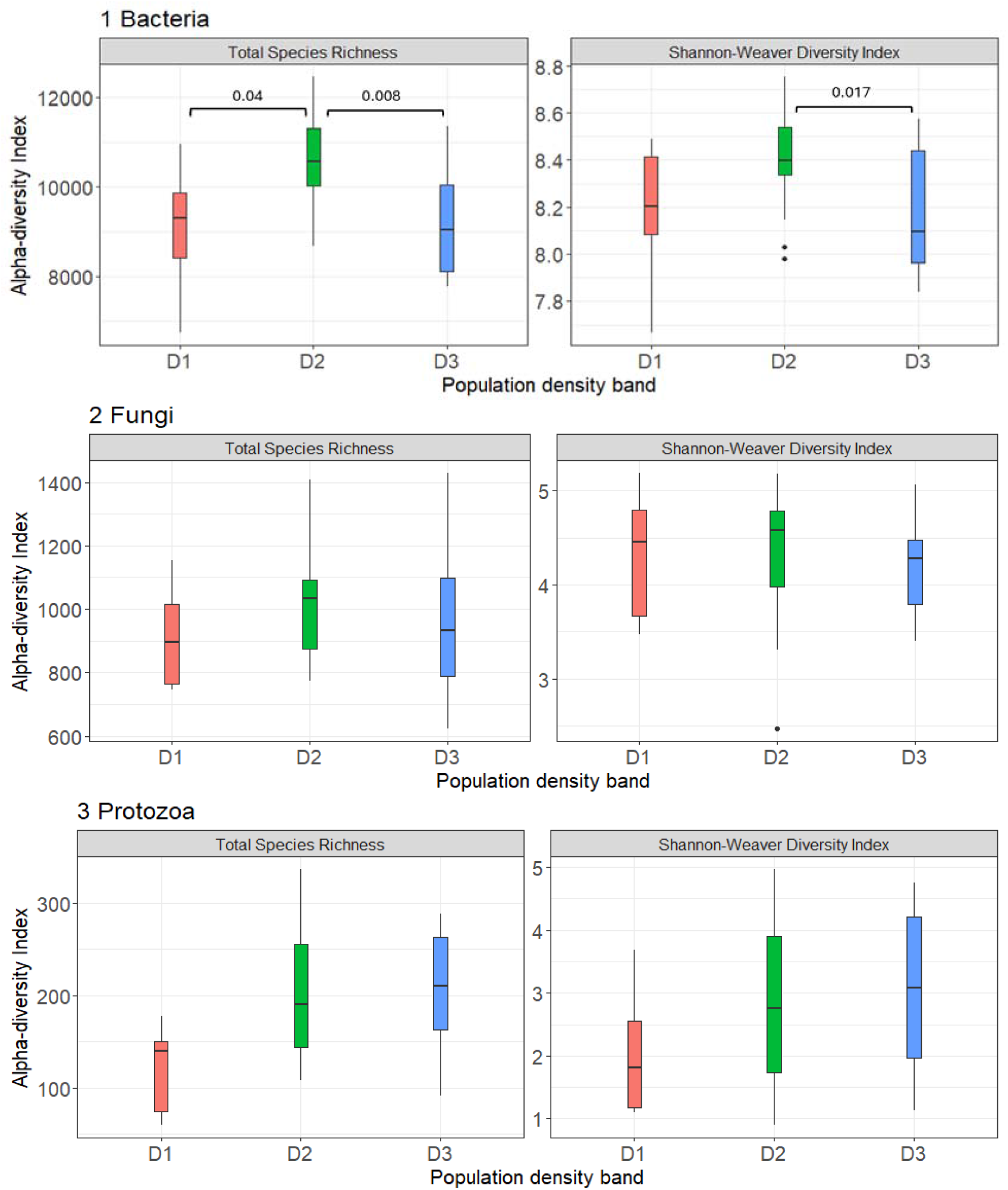
Boxplots showing the distribution of soil microbial alpha-diversity indices across population density bands. Displayed are the alpha-diversity of (a) bacteria, (b) fungi, and (c) protozoa zOTU’s across population density bands: D1 = 10 – 837 population/km^2^ (red), D2 = 1035 – 2203 population/km^2^ (green), and D3 = 2500 – 3752 population/km^2^ (blue). Panels show alpha-diversity calculated using Total Species Richness and Shannon-Weaver Diversity Index. Boxplots represent group median with interquartile range. Outliers are indicated as dots. Statistical significance between groups is shown (p < 0.05).

### 3.2 Beta-diversity

Beta-diversity did not differ between the three population density bands for soil bacteria (PERMANOVA, F=1.2, R^2^=0.057, p=0.091) or fungi (PERMANOVA, F=1.1, R2=0.054, p=0.17). Soil protozoan communities showed a significant PERMANOVA result (PERMANOVA, F=1.2, R^2^=0.061, p=0.044), however after Bonferroni-correction, pairwise comparisons revealed no differences between population density bands. Principal component analysis was used to visualise the PERMANOVA results and was able to explain 18.4% (bacteria), 14.2% (fungi), and 16.8% (protozoa) of the total variation in the data for microbiome analyses. Population density as an ordination vector was not significant in explaining the variation between sites (Figure 3, Table S7).

**Figure 3.**
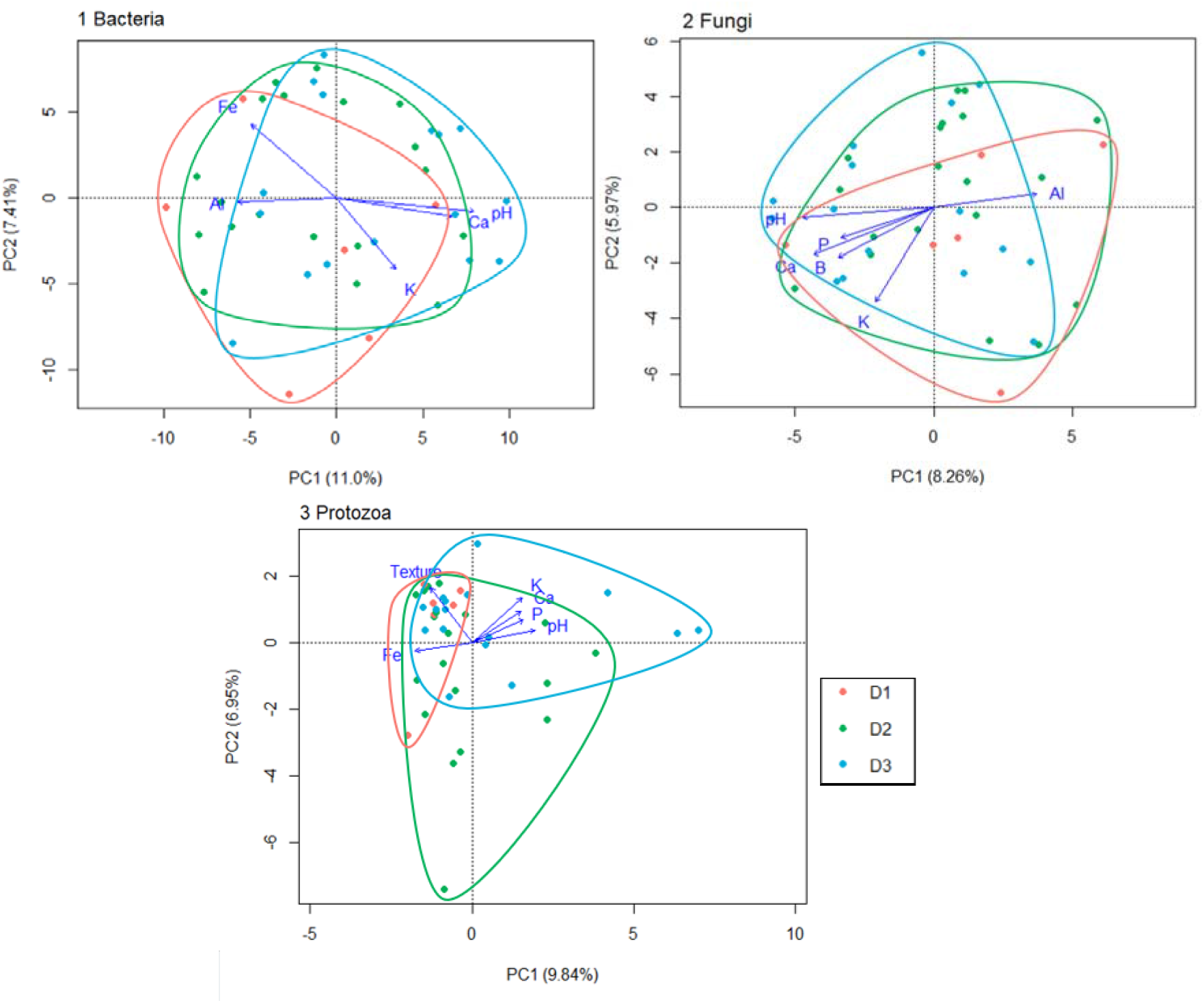
Principal component analysis (PCA), based on Aitchison distance, of soil microbial community composition across population density bands in Hobart. PCA plot showing dissimilarity between centred log-ratio transformed (1) bacterial, (2) fungal, and (3) protozoan zOTU community composition across density bands: D1 = 10 – 837 population/km^2^ (red), D2 = 1035 – 2203 population/km^2^ (green), and D3 = 2500 – 3752 population/km^2^ (blue). Displayed are sites as dots, coloured by population density band. The primary and secondary axes, PC1 and PC2, are plotted and the percentage variation explained are indicated in the axes parentheses. Blue arrows indicate significant environmental predictor variables to the ordination variation (Bonferroni-adjusted threshold p < 0.05). Vector lengths are scaled to the strength of the association (see Table S7). Fe=iron, Al=aluminium, Ca=calcium, K=potassium, P=phosphorus, B=boron.

### 3.3 Community Composition

All three population density bands were dominated by the same top phyla (Figure 4). Bacterial communities were dominated by Actinobacteria, with Solirubrobacterales and Nocardioidaceae appearing across most sites, and Proteobacteria, primarily from the Xanthobacteraceae family.

**Figure 4.**
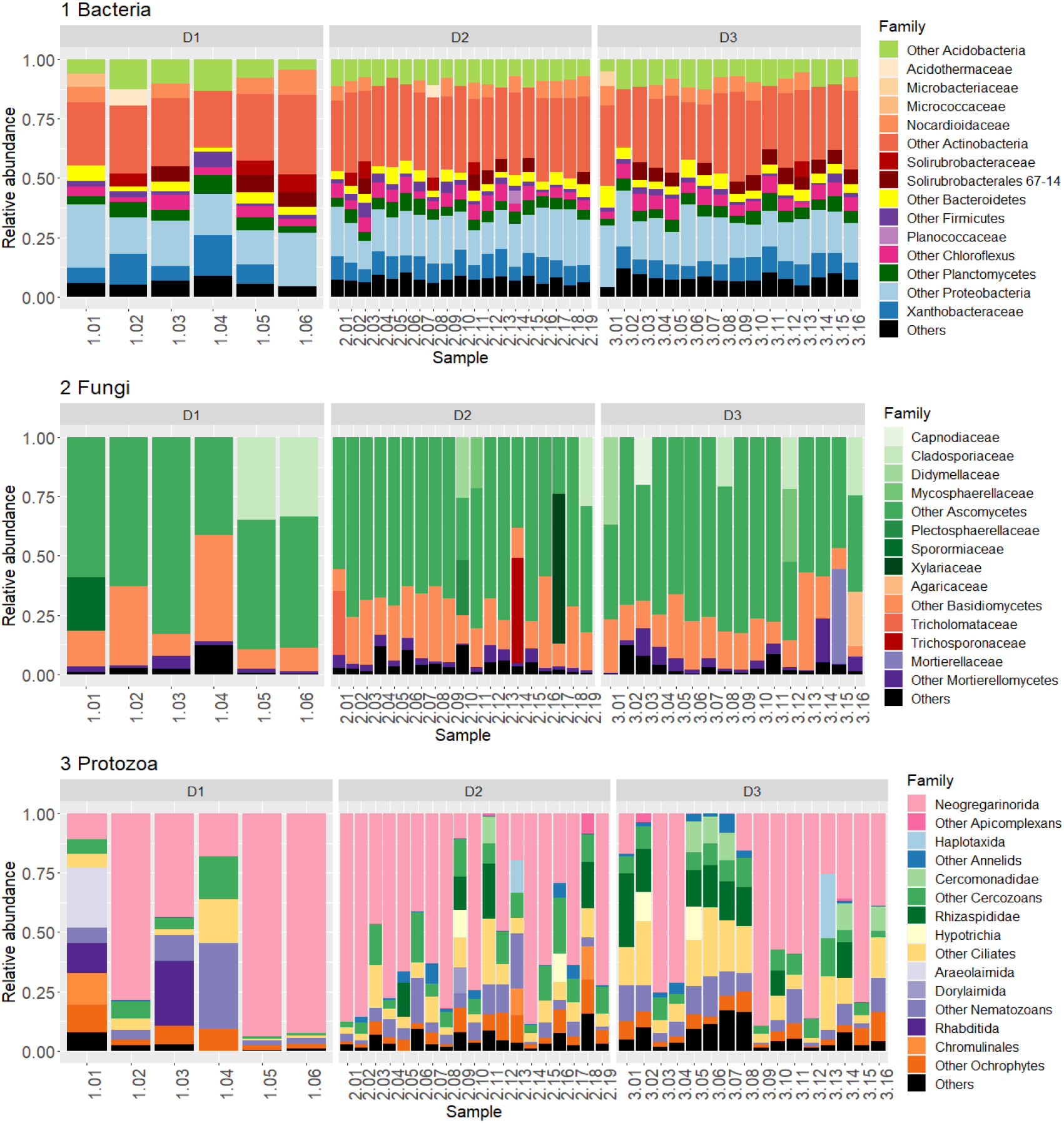
Soil microbial community composition at the phylum/family level for each sampling site. Panels show (1) bacterial, (2) fungal, and (3) protozoan communities for sampling locations across population density bands: D1 = 10 – 837 population/km^2^, D2 = 1035 – 2203 population/km^2^, and D3 = 2500 – 3752 population/km^2^. Displayed are relative abundances (%) of the top families (>5% bacteria, >20% fungi, 10% protozoa), grouped by colour into their respective phylum, for the -16S rRNA, -ITS, and -18S rRNA amplicons. All phylum not belonging to a top family group are grouped into “Others”. For bacteria, family with a relative abundance <5% are grouped into “Others/ Other phyla”. For fungi, family with a relative abundance 20% are grouped into “Others/ Other phyla”. For protozoa, family with a relative abundance <10% are grouped into “Others/ Other phyla”.

Fungal communities were dominated by Ascomycetes and Basidiomycetes. Most Ascomycetes families within each site were represented in relative abundances lower than 20%, so could not be displayed clearly on Figure 4. However, some sites showed high dominance of certain family groups, like Xylariaceae and Cladosporiaceae, without a clear trend with population density band.

Protozoan communities were mostly dominated by organisms belonging to the Apicomplexans phylum, Neogregarinorida order, as well as Nematozoans, Cercozoans and Ciliates phylum. However, there was high between-site variation in community composition within all the population density band groups. Sites were either (1) quite diverse in phyla and associated family representation, with most of the top phyla listed above appearing within a site in high relative abundance (> 10%). Or, (2) sites were dominated primarily by neogregarines, with many sites representing this family with 50% relative abundance or greater and little representation of the remaining top phylum. This trend was observed across all sites, irrespective of population density band.

### 3.4 Soil physicochemical influence on soil communities

The three population density bands had the same constitution of soil physicochemical variables, except for soil potassium (Table S6). Soil potassium concentrations were greater in the high population density band, compared to the middle density band, D2 (Table S6).

#### 3.4.1 Soil physicochemistry and its association with alpha-diversity indices

We found no correlations of soil physicochemical variables to alpha-diversity indices for bacterial and fungal communities (Figure S3). Protozoan alpha-diversity in low population density sites was inversely proportional to soil manganese concentrations only. There were no correlations in protozoan alpha-diversity to soil variables across the middle (D2) and high population density (D3) groups (Figure S3).

#### 3.4.2 Soil physicochemistry and its association with beta-diversity

Several soil variables were significantly correlated with the variation between sites of the principal component ordination computed (Figure 3, Table S7). Based on the *envfit* calculated correlations, pH exhibited the strongest correlation to the variation between bacterial communities (R^2^ = 0.74, p = 0.002). Across all three microbial domains, similar soil variables were associated with variation in community composition in weak and moderate correlations, and include pH, calcium and potassium; while phosphorus, iron, aluminium, soil texture and boron exhibited kingdom-specific associations (see Table S7).

## 4 Discussion

Using a multi-kingdom approach in a regional city, we tested whether population density is associated with variation in urban soil microbial communities. Population density was associated with bacterial alpha-diversity only, which followed a non-linear, inverse U-shaped pattern, with highest diversity at intermediate densities. In contrast, fungal communities showed no response to population density across any diversity or compositional metric, and microbial community composition (beta-diversity) did not differ across density bands for any group. Protozoan communities varied markedly among sites, but this variation was not structured by population density, suggesting the influence of other, unmeasured urban-related drivers. Together, these findings provide clear evidence against one-size-fits-all assumptions about urban impacts on soil microbiomes and indicate that both microbial kingdom and urban context mediate how soil communities respond to urbanisation.

### 4.1 Bacterial diversity across population density bands

Soil bacterial communities were the only microbial group to exhibit a significant response to population density, with alpha-diversity following a non-linear, inverse U-shaped relationship. Alpha-diversity peaked at intermediate population densities before declining at the highest density sites, suggesting that moderate levels of urbanisation may create heterogeneous soil conditions capable of supporting a wider range and abundance of bacterial taxa. In contrast, both low- and high-population density areas may impose stronger environmental constraints, resulting in reduced bacterial diversity. Although richness and evenness varied across population density bands, bacterial community composition remained comparatively stable, suggesting that population density may influence bacterial diversity through changes in taxa abundance and persistence rather than broad community restructuring.

These findings align with the Intermediate Disturbance Hypothesis (IDH), which predicts that intermediate disturbance can promote diversity by increasing niche availability and reducing competitive exclusion, while avoiding the extreme stress associated with heavily disturbed systems (Connell, 1978). The urban environment is full of human-driven disturbances at an intermediate level, including foot traffic, lawn mowing, irrigation and pet activity, all of which may periodically disrupt soil structure, and introduce organic inputs like leaf litter, grass clippings, animal scat, water, and fertiliser. Collectively, these processes may increase microhabitat variability and resource availability, creating conditions that can be exploited by a wider range of opportunistic bacterial taxa.

The increase in bacterial diversity observed at intermediate population densities is consistent with previous studies demonstrating that urban environments can support greater bacterial diversity compared with less urbanised areas (Delgado-Baquerizo et al., 2021; Gao et al., 2023; Li et al., 2023; Ramirez et al., 2014; Wang et al., 2018). Urban landscapes can act as local hotspots of soil microbial diversity by creating heterogeneous habitats with varied environmental conditions and resource inputs (Delgado-Baquerizo et al., 2021; Ramirez et al., 2014). In large urban centres, human population density as a proxy for urban intensity has been positively associated with bacterial diversity in urban greenspaces (Wang et al., 2018), while studies across urbanisation gradients have similarly reported increases in bacterial diversity with increasing urban intensity (Gao et al., 2023; Li et al., 2023). At a smaller spatial scale, Grierson et al. (2023) found that soil bacterial alpha-diversity was lowest in public greenspaces experiencing lower levels of human activity in comparison to areas experiencing higher levels of human activity as well as residential backyards across a regional city, further suggesting that moderate anthropogenic disturbance may enhance microbial diversity. Collectively, these studies indicate that urban environments can create a mosaic of altered soil conditions, disturbance regimes, and resource inputs that may increase habitat complexity and provide opportunities for coexistence among bacterial taxa. However, many previous studies have also reported concurrent shifts in bacterial community composition, unlike the present study where increased bacterial diversity occurred without substantial turnover in bacterial assemblages. This indicates that bacterial diversity can vary across the urban gradient without necessarily being accompanied by substantial changes in overall community composition.

### 4.2 Why fungi and protozoa did not respond to population density

In contrast to bacteria, fungal communities did not vary significantly in either alpha- or beta-diversity across population density bands, suggesting that broad measures of urbanisation may have limited capacity to influence fungal diversity. This is consistent with the ecology of soil fungi, which are closely associated with vegetation through plant–fungal interactions and the decomposition of organic matter (Frac et al., 2018). Accordingly, previous studies along urban gradients have shown that fungal communities are often structured more strongly by vegetation characteristics, local habitat conditions, and environmental variables than by broad urbanisation metrics (Hui et al., 2017; Tan et al., 2019). Although some studies have reported changes in fungal diversity or community composition with increasing urbanisation (Abrego et al., 2020; Rusterholz & Baur, 2023), these responses appear less consistent than those observed for bacteria and may reflect differences in vegetation, urban context, and the fungal functional groups considered.

Protozoan communities showed only weak associations with population density, with no statistically significant changes in alpha-diversity and only a weak association between community composition and population density. Although differences in community composition were detected among sampling sites, these differences were not clearly structured by population density, suggesting that local site characteristics or other environmental factors not captured by population density may exert a stronger influence on protozoan community assembly. This interpretation is consistent with emerging evidence that soil protists are primarily structured by fine-scale environmental conditions and trophic interactions. For example, Shangguan et al. (2024) found that the drivers of protist communities differed among functional groups, with bacterivorous protists responding primarily to bacterial communities, while phototrophic and parasitic protists were more strongly associated with soil nutrient availability and abiotic conditions. Likewise, Fiore-Donno et al. (2019) demonstrated that soil protist communities were influenced by local edaphic properties, including soil moisture, clay content, and nutrient availability, highlighting the importance of microsite environmental heterogeneity in structuring protist assemblages. Although relatively few studies have examined soil protists across urbanisation gradients, the available evidence, together with the findings of the present study, suggests that microbial responses to urbanisation are taxon-specific and population density is likely too coarse a proxy to capture the fine-scale environmental and biological processes governing fungal and protozoan community assembly.

### 4.3 Composition across population density bands

Despite differences in bacterial alpha-diversity, microbial community composition remained consistent across the population density gradient. Several soil physicochemical variables were significantly associated with variation in microbial community composition, although population density itself was not. These findings suggest that local soil conditions influence microbial community composition more than population density alone.

In the present study, population density was not associated with measured soil physicochemical variables, suggesting that the environmental changes accompanying urbanisation were insufficient to generate strong environmental filtering capable of excluding or replacing bacterial taxa. Instead, urbanisation may have altered the relative abundance and persistence of taxa already present within a shared regional species pool. Comparable results were reported within the medium-sized regional city of Dijon, France, where microbial community composition showed only modest differentiation across urban environments (Christel et al., 2023).

At broader spatial scales, emerging evidence indicates that urbanisation can increase local microbial diversity while simultaneously driving compositional homogenisation. A global assessment of urban greenspaces demonstrated that urban soils supported high bacterial and protist diversity despite microbial communities becoming increasingly similar across cities worldwide (Delgado-Baquerizo et al., 2021), a pattern consistent with Zhang et al. (2025), who found increased homogenisation of bacterial communities across urban soils. Thus, greater local microbial diversity does not necessarily mean that urban soils contain increasingly distinct microbial communities, as the additional diversity may include taxa that are widely shared across urban environments. Urban green spaces may therefore support diverse microbial communities locally, while exposure across different cities may increasingly involve similar suites of microbial taxa. Whether this increasing homogenisation has consequences for human health or the ecosystem functions provided by urban soil microbiomes remains an important question

### 4.4 City size matters: implications of studying a regional city

Current understanding of urban soil microbiomes is heavily derived from studies conducted in large, rapidly urbanising metropolitan regions. While these systems provide valuable insight into the ecological consequences of intensive urbanisation, they may not adequately represent the diversity of urban environments globally. Small- and medium-sized cities accommodate a substantial proportion of the world’s urban population (United Nations, 2025) and account for much of ongoing urban growth yet remain comparatively understudied in urban biodiversity research (Kendal et al., 2020). Urbanisation within these cities is often more heterogeneous, less intensive, and characterised by a mosaic of residential, greenspace and remnant natural habitats, potentially producing different ecological responses from those observed in megacities.

Urbanisation should therefore be viewed as a continuous ecological process rather than a binary urban–non-urban state. The environmental characteristics commonly associated with highly urbanised metropolitan regions—including extensive impervious surface cover, pronounced urban heat island effects, elevated pollution and anthropogenic landscape homogenisation – can vary substantially with the size and intensity of urban development (Chien, 2025; Wei et al., 2024). These characteristics are unlikely to represent universal features of all urban environments but rather emerge progressively as cities develop. Consequently, microbial responses to urbanisation may also vary across different stages of urban development. Future studies spanning cities of different sizes and developmental trajectories will be essential to identify drivers of soil microbiome diversity and to improve the generalisability of urban soil microbiome research.

### 4.5 Limits of population density and directions for future urban microbiome research

Overall, our study contributes to a growing body of evidence emphasising the limitations of population density as a means by which to understand (or predict) the impacts of urbanisation on soil microbiomes. Clearly, although not without its utility, population density remains a coarse metric for urbanisation, which is not always of prime relevance to understanding soil microbiome variation. Even where associations between microbiome patterns and population density are observed, it is likely that population density is acting as an ‘umbrella’ proxy for a complex and context-dependent mixture of underlying drivers. We argue that studies should focus more directly on measuring associations between microbiome communities and such drivers, in order to facilitate a more nuanced understanding.

Specifically, we argue for a multi-metric urbanisation framework, incorporating multiple factors such as land use history, land & soil management practices, vegetation history and chemical inputs, if available. All of these factors have been associated with soil microbiome diversity and/or composition in diverse contexts (Christel et al., 2023; Thompson et al., 2023) and have clear mechanistic pathways linking them to microbiome patterns. We further argue that studies should assess relationships between urbanisation and soil microbiomes in a way that accounts for the inherently high inter- (and intra-) site variability in soil microbiome diversity and structure (Delgado-Baquerizo et al., 2021; Yan et al., 2016). Our study also underlines the value of multi-kingdom studies, rather than relying on soil bacterial studies alone. Multi-kingdom designs, although more costly, are clearly necessary to understand how different components of soil microbiomes respond to different urban-related environmental drivers.

A key limitation of our study is that we did not have access to longitudinal data - in particular, pre-urbanisation data - with which to assess the impacts of urban-related land use change over time. Given the increasing evidence that land-use histories and land-use change trajectories can influence urban microbiome patterns (Louisson et al., 2023; Thompson et al., 2023), we highlight the value of longitudinal studies for deepening our understanding of the nuances of urbanisation-soil microbiome relationships.

## 5 Conclusion

This study demonstrates that population density is a selective and context-dependent driver of urban soil microbiomes rather than a universal indicator of urban impact. In a regional city, population density was associated with bacterial alpha-diversity only, following a non-linear pattern, while fungal and protozoan communities showed no consistent response. Community composition across all microbial groups was structured more strongly by soil physicochemical properties than by population density. These results highlight that microbial responses to urbanisation are kingdom-specific, therefore extrapolating conclusions from bacteria alone or from highly urbanised megacity contexts is inappropriate.

By focusing on a small, regional city, reflective of where most urban residents reside globally, this work challenges one-size-fits-all assumptions about urban effects on soil microbiomes and underscores the importance of urban context and scale in environmental microbiology. More broadly, our findings emphasise the need for multi-kingdom, multi-metric approaches that integrate human pressures, soil properties, and land-use dynamics to better understand how urbanisation reshapes soil ecosystems. Such approaches are essential for informing urban planning, biodiversity conservation, and the management of urban environments in ways that support resilient soil microbiomes, and the ecosystem functions they underpin.

