## Supplementary material for "Population density is a selective driver of urban soil microbiomes: evidence from bacteria, fungi and protozoan communities in a regional city"

Corresponding author: Jessica Grierson

**Supplementary Material**

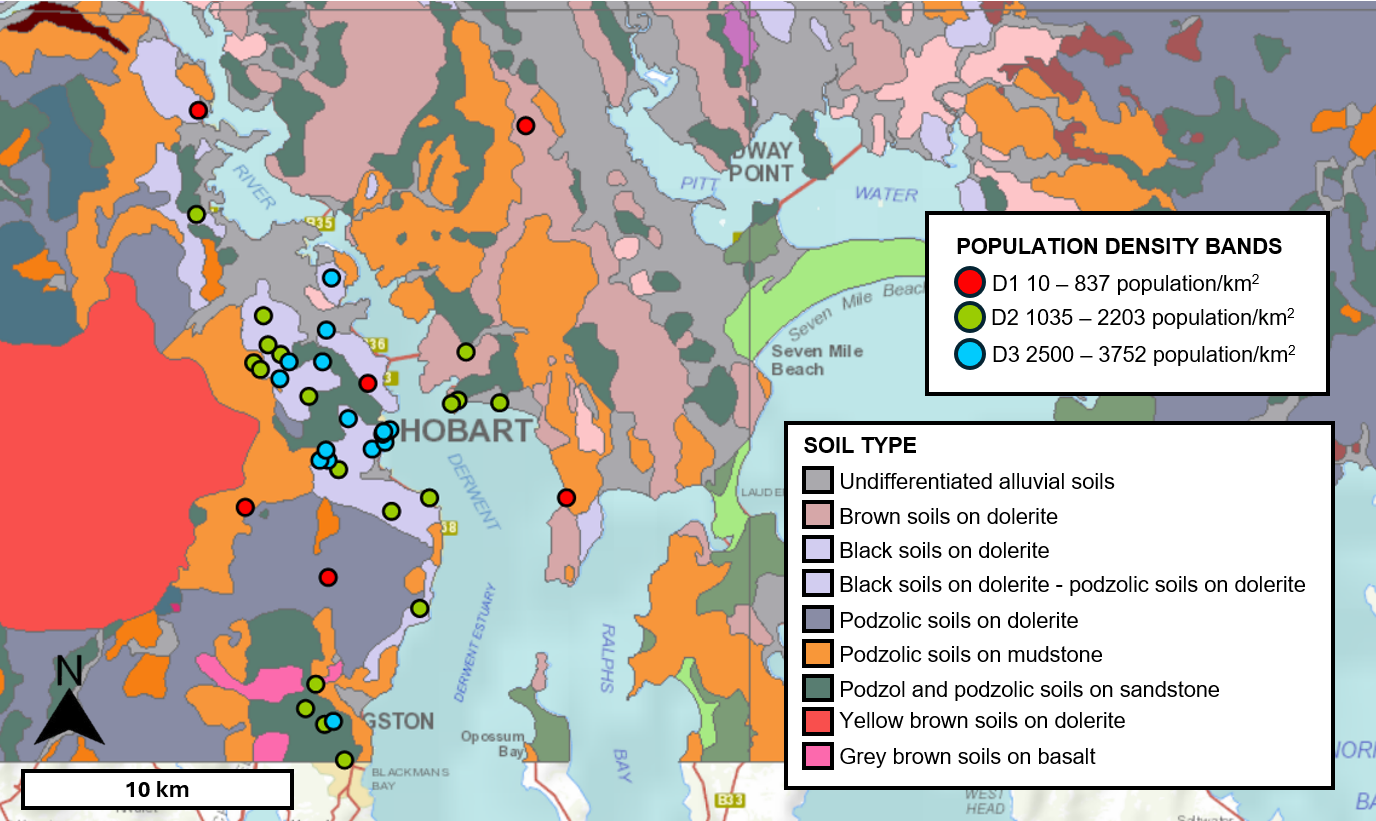

**Figure S1 Map of population density band sampling locations on soil types in Hobart, Australia**. Shown are soil sampling locations across population density bands: D1 (10 – 837 population/km^2^; red), D2 (1035 – 2203 population/km^2^; green) and D3 (2500 – 3752 population/km^2^; blue). Background shading shows soil types across the greater Hobart region. Information was sourced from ListMAP (Land Tasmania, 2022) based on SA1 and DPIPWE Reconnaissance Soil Map Series of Tasmania (Spanswick & Kidd, 2000)

**Table S1 Count of soil samples collected along different soil types in Hobart, Australia.**

| SOIL TYPE | D1 | D2 | D3 |
| --- | --- | --- | --- |
| Undifferentiated alluvial soils (A) |  | 1 | 2 |
| Brown soils on dolerite (Bd1) | 1 | 1 |  |
| Black soils on dolerite (Bld1) | 2 | 9 | 11 |
| Black soils on dolerite-podzolic soils on dolerite (Bld1-Pd1) |  |  | 1 |
| Podzolic soils on dolerite (Pd) | 1 |  |  |
| Podzolic soils on mudstone (Pm) | 2 | 4 |  |
| Podzol and podzolic soils on sandstone (Pss) |  | 4 | 2 |
| Total | 6 | 19 | 16 |

**Table S2 Soil physical and chemical properties measured for each soil sample**. Table adapted from Bissett et al. (2016).

| *Soil chemical properties* | | |
| --- | --- | --- |
| Ammonium nitrogen | Conductivity | Aluminium |
| Nitrate nitrogen | pH | Calcium |
| Phosphorus (Colwell) | Copper | Magnesium |
| Potassium (Colwell) | Iron | Potassium |
| Sulphur | Manganese | Sodium |
| Organic carbon | Zinc | Boron |
| *Soil physical properties* | | |
| Texture | Elevation |  |

**Figure S2 Rarefaction curves per site for soil bacteria, fungi, and protozoa.**

1 Bacteria

2 Fungi

3 Protozoa

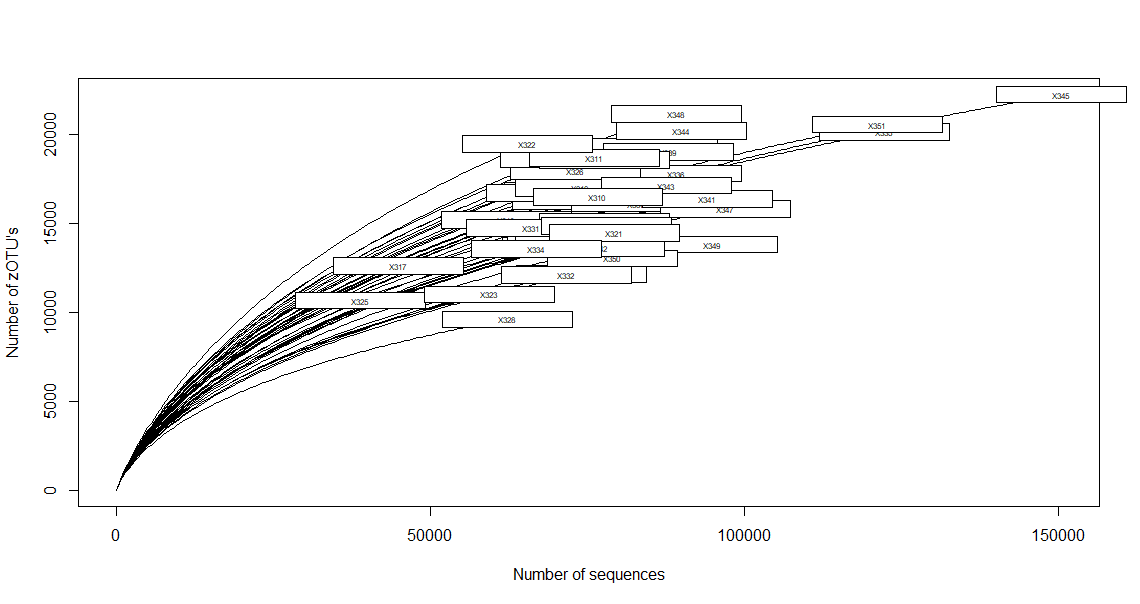

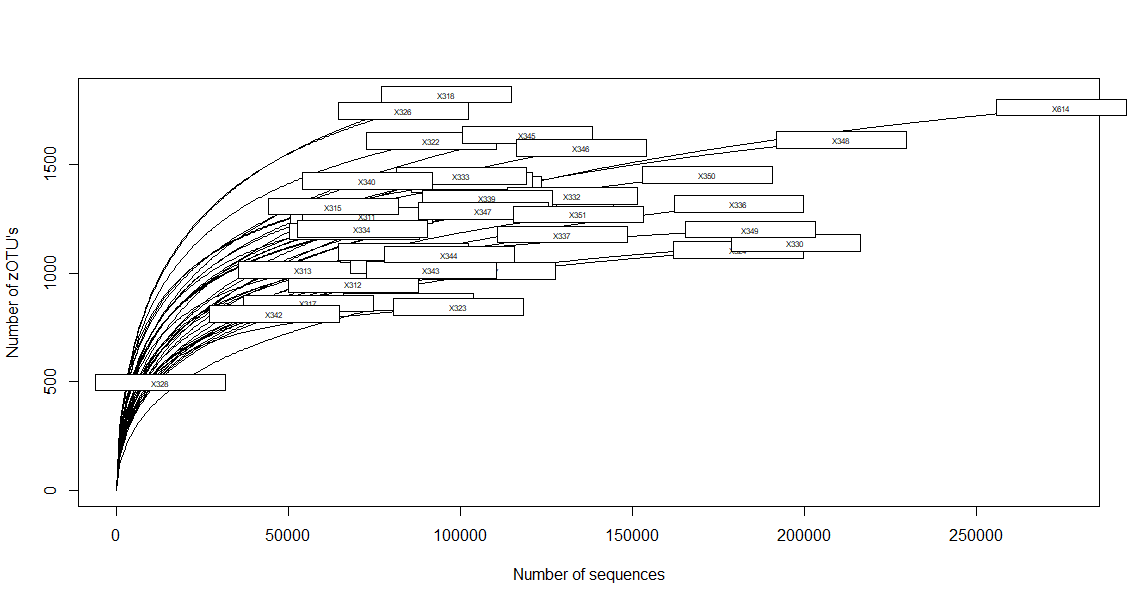

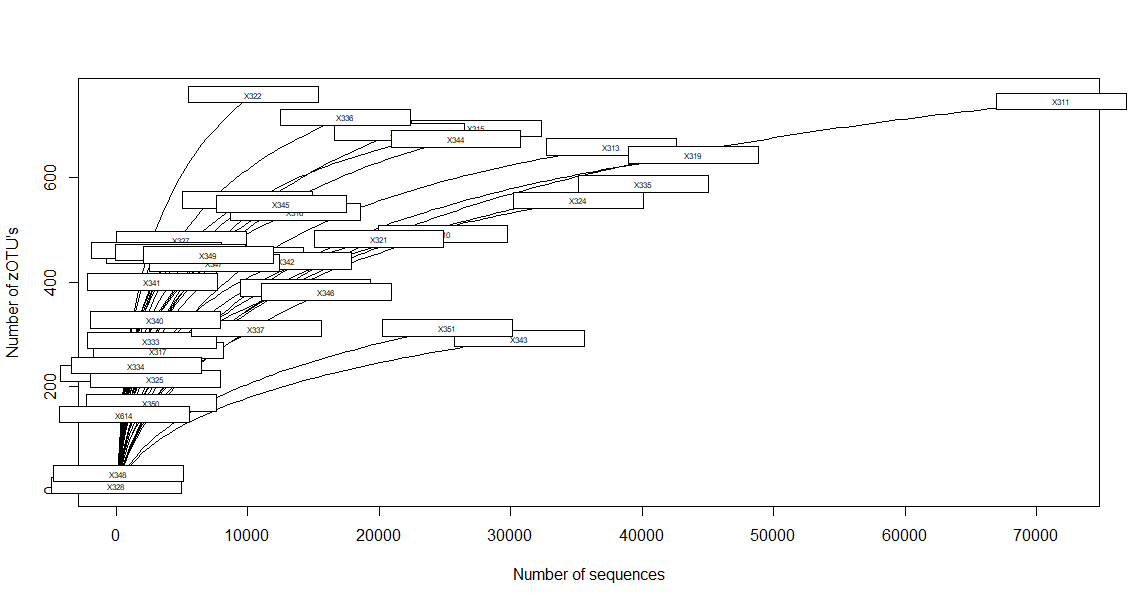

**Table S3 Significance values for alpha-diversity normality tests across population density bands.** Shapiro-Wilk test was used to check for normal distribution, and Bartlett’s test was used to test for equal variance for bacterial, fungal, and protozoan alpha-diversity indices. Population density bands: D1 = 10 – 837 population/km^2^, D2 = 1035 – 2203 population/km^2^, and D3 = 2500 – 3752 population/km^2^.

|  | | | Total Species Richness | Shannon-Weaver Diversity Index |
| --- | --- | --- | --- | --- |
| Bacteria | *Shapiro-Wilk* | D1 | 0.95 | 0.46 |
|  |  | D2 | 0.77 | 0.60 |
|  |  | D3 | 0.11 | 0.056 |
|  | *Bartlett’s* | | 0.65 | 0.44 |
| Fungi | *Shapiro-Wilk* | D1 | 0.57 | 0.60 |
|  |  | D2 | 0.29 | 0.059 |
|  |  | D3 | 0.68 | 0.46 |
|  | *Bartlett’s* | | 0.85 | 0.46 |
| Protozoa | *Shapiro-Wilk* | D1 | 0.44 | 0.44 |
|  |  | D2 | 0.43 | 0.37 |
|  |  | D3 | 0.18 | 0.15 |
|  | *Bartlett’s* | | 0.77 | 0.88 |

**Table S4 Significance values for differences in soil microbial alpha-diversity indices across population density bands.** One-way analysis of variance (ANOVA) was used to compare across all three sampling locations for bacteria, fungi, and protozoa. Tukey honest significance test (Tukey HSD) was used to test differences between sampling locations and bonferroni adjusted p-values are reported. Statistical significance is marked in bold (corrected p-value threshold < 0.05). Population density bands: D1 = 10 – 837 population/km^2^, D2 = 1035 – 2203 population/km^2^, and D3 = 2500 – 3752 population/km^2^.

|  | Total Species Richness | | | Shannon-Weaver Diversity Index | | |
| --- | --- | --- | --- | --- | --- | --- |
| *Sampling location* | Bacteria | Fungi | Protozoa | Bacteria | Fungi | Protozoa |
| ANOVA | **0.004** | 0.46 | 0.059 | **0.014** | 0.82 | 0.33 |
| D1 – D2 | **0.039** | 0.58 | 0.068 | 0.13 | 0.99 | 0.37 |
| D1 – D3 | 0.97 | 0.95 | 0.064 | 0.99 | 0.91 | 0.32 |
| D2 – D3 | **0.008** | 0.56 | 0.99 | **0.017** | 0.83 | 0.98 |

**Table S5 Dispersion (*betadisper*) and pairwise PERMANOVA (*adonis2*) group model results for differences in beta-diversity, based on Aitchison distance, of soil microbial communities across population density bands.** *betadisper* was used to test for homogeneity of variance. Permutational multivariate analysis of variance (PERMANOVA) was used to compare across all three population density bands for bacteria, fungi, and protozoa using *adonis2*. Tukey honest significance test (Tukey HSD) was used to test differences between sampling locations and bonferroni-adjusted p-values are reported. Statistical significance is marked in bold (corrected p-value threshold < 0.05). Population density bands: D1 = 10 – 837 population/km^2^, D2 = 1035 – 2203 population/km^2^, and D3 = 2500 – 3752 population/km^2^.

|  |  | | F-model | R2 | P-value |
| --- | --- | --- | --- | --- | --- |
| Bacteria | *betadisper* | | 0.73 | - | 0.49 |
|  | *adonis2* | | 1.2 | 0.057 | 0.091 |
|  |  | *D1 – D2* | 1.1 | 0.044 | 0.88 |
|  |  | *D1 – D3* | 1.2 | 0.057 | 0.41 |
|  |  | *D2 – D3* | 1.2 | 0.035 | 0.30 |
| Fungi | *betadisper* | | 0.41 | - | 0.67 |
|  | *adonis2* | | 1.1 | 0.054 | 0.17 |
|  |  | *D1 – D2* | 1.0 | 0.042 | 1.0 |
|  |  | *D1 – D3* | 1.1 | 0.051 | 0.73 |
|  |  | *D2 – D3* | 1.1 | 0.033 | 0.39 |
| Protozoa | *betadisper* | | 0.79 | - | 0.46 |
|  | *adonis2* | | 1.2 | 0.061 | **0.044** |
|  |  | *D1 – D2* | 1.1 | 0.047 | 0.68 |
|  |  | *D1 – D3* | 1.5 | 0.068 | 0.13 |
|  |  | *D2 – D3* | 1.2 | 0.034 | 0.31 |

**Table S6 Soil physicochemical variables and pairwise comparisons for differences across population density bands.** Shown is the mean ± standard error of soil variables across population density bands: D1 (10 – 837 population/km^2^), D2 (1035 – 2203 population/km^2^), and D3 (2500 – 3752 population/km^2^). Kruskal-Wallis rank sum test was used to test for significant differences between population density bands, and a Wilcoxon signed-rank test was used for pairwise comparisons between sampling locations. Pairwise comparison results are listed for the statistically significant Kruskal-Wallis results, and Bonferroni adjusted p-values are reported in parentheses (p < 0.05). Texture category code: 1=Sand, 1.5=Sand/Loam, 2=Loam, 2.5=Loam/Clay, 3=Clay, 3.5=Very heavy clay.

| Soil physicochemical variables | Population density band | | | Pairwise comparison |
| --- | --- | --- | --- | --- |
|  | D1 | D2 | D3 |  |
| Potassium (mg/Kg) | 369 ± 90.7 | 302 ± 40.5 | 466 ± 47.6 | D2 – D3  (0.028) |
| Elevation (m) | 94.6 ± 36.9 | 69.0 ± 10.4 | 57.9 ± 10.2 |  |
| Texture (category code) | 2.06 ± 0.11 | 1.91 ± 0.051 | 1.89 ± 0.046 |  |
| Fine sand (%) | 30.7 ± 0.94 | 27.6 ± 1.39 | 29.3 ± 0.93 |  |
| Sand (%) | 63.9 ± 3.84 | 70.8 ± 2.25 | 66.3 ± 1.49 |  |
| Silt (%) | 12.0 ± 1.56 | 9.66 ± 1.30 | 11.1 ± 0.68 |  |
| Clay (%) | 24.1 ± 2.41 | 19.6 ± 1.36 | 22.7 ± 1.19 |  |
| Ammonium nitrogen (mg/Kg) | 6.06 ± 2.58 | 17.2 ± 3.05 | 20.2 ± 6.90 |  |
| Nitrate nitrogen (mg/Kg) | 10.1 ± 3.53 | 12.8 ± 3.17 | 13.5 ± 3.04 |  |
| Phosphorus (mg/Kg) | 60.5 ± 22.9 | 80.1 ± 9.68 | 83.4 ± 10.4 |  |
| Sulphur (mg/Kg) | 7.84 ± 0.86 | 11.7 ± 1.13 | 11.2 ± 1.27 |  |
| Organic carbon (%) | 3.56 ± 0.34 | 3.62 ± 0.18 | 3.77 ± 0.18 |  |
| Conductivity (dS/m) | 0.12 ± 0.018 | 0.12 ± 0.008 | 0.15 ± 0.011 |  |
| pH | 5.60 ± 0.25 | 5.44 ± 0.11 | 5.58 ± 0.13 |  |
| Copper (mg/Kg) | 2.97 ± 0.85 | 3.25 ± 1.15 | 3.89 ± 0.62 |  |
| Iron (mg/Kg) | 126 ± 17.3 | 181 ± 16.3 | 169 ± 16.8 |  |
| Manganese (mg/Kg) | 19.4 ± 5.31 | 18.0 ± 3.04 | 19.9 ± 2.93 |  |
| Zinc (mg/Kg) | 48.2 ± 14.4 | 46.8 ± 10.2 | 63.3 ± 11.0 |  |
| Aluminium (meq/100g) | 0.15 ± 0.054 | 0.11 ± 0.022 | 0.081 ± 0.007 |  |
| Calcium (meq/100g) | 14.8 ± 2.26 | 13.6 ± 1.15 | 17.4 ± 1.09 |  |
| Magnesium (meq/100g) | 4.22 ± 0.87 | 3.35 ± 0.40 | 4.16 ± 0.39 |  |
| Sodium (meq/100g) | 0.37 ± 0.095 | 0.28 ± 0.029 | 0.31 ± 0.037 |  |
| Boron (mg/Kg) | 1.26 ± 0.24 | 1.01 ± 0.093 | 1.15 ± 0.11 |  |

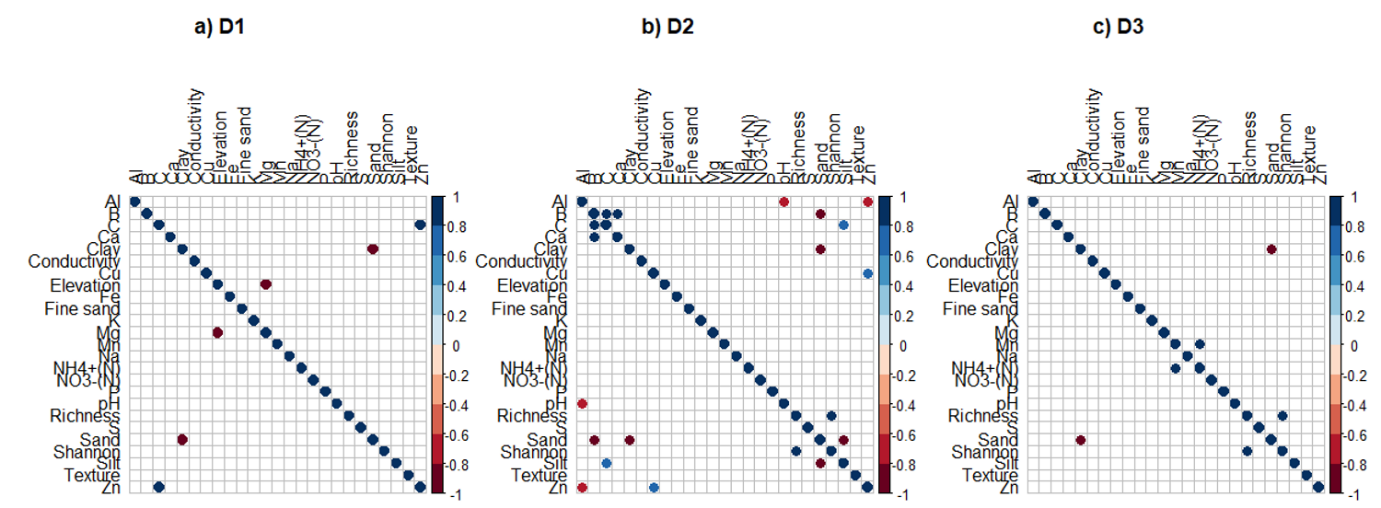

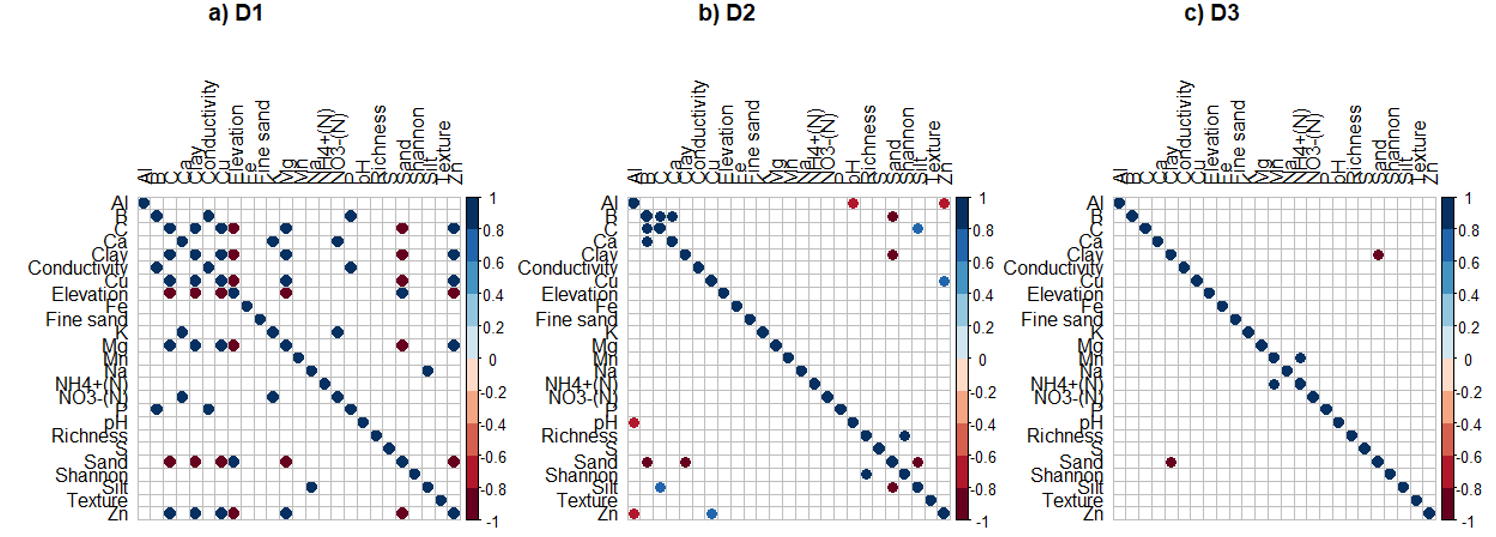

**1 Bacteria**

**2 Fungi**

**3 Protozoa**

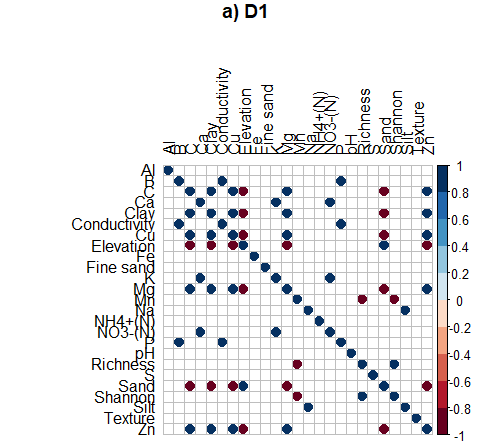

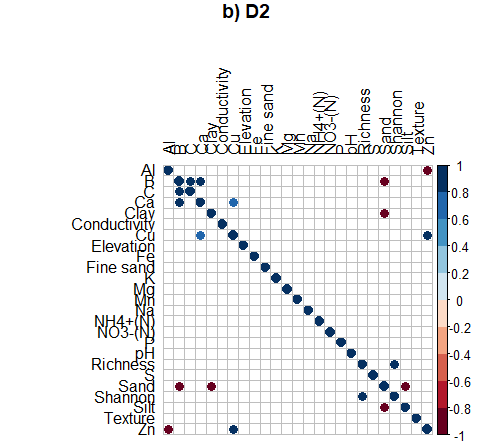

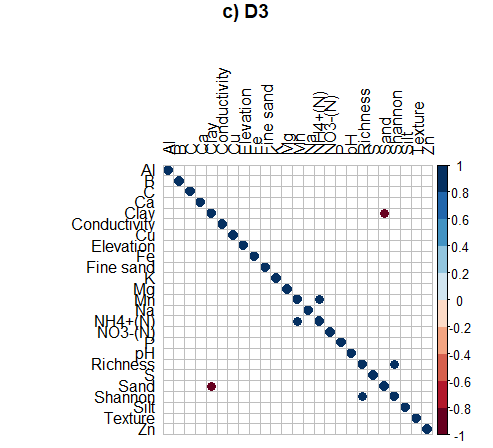

**Figure S3 Spearman’s rank-order correlation matrix between alpha-diversity indices and soil contextual variables within population density bands for major taxonomic groups.** Displayed are statistically significant correlations between environmental predictor variables and alpha-diversity indices as circles (Holm-adjusted

p < 0.05) for soil (1) bacteria, (2) fungi and (3) protozoa in population density bands (a) D1 (10 – 837 population/km^2^), (b) D2 (1035 – 2203 population/km^2^), and (c) D3 (2500 – 3752 population/km^2^). Magnitude and direction of correlation coefficient depicted by colour of circle. Larger circles indicate a stronger correlation between variables. Al=aluminium, B=boron, C=organic carbon, Ca=calcium, Cu=copper, Fe=iron, K=potassium (Colwell), Mg=magnesium, Mn=manganese, Na=sodium, NH4+(N)=ammonium nitrogen, NO3-(N)=nitrate nitrogen, P=phosphorus (Colwell), Richness=Total Species Richness, S=sulphur, Shannon=Shannon-Weaver Diversity Index, Zn=zinc.

**Table S7 Environmental predictor variables predicting soil microbial community compositions with associated *envfit* R2 and Bonferroni-adjusted p-values for bacterial, fungal, and protozoan population density band PCA in Figure 3.2.** Statistical significance is marked in bold (Bonferroni-adjusted threshold

p < 0.05). Texture category code: 1=Sand, 1.5=Sand/Loam, 2=Loam, 2.5=Loam/Clay, 3=Clay, 3.5=Very heavy clay.

| Environmental predictor variables | Bacteria | | Fungi | | Protozoa | |
| --- | --- | --- | --- | --- | --- | --- |
|  | R^2^ | P-value | R^2^ | P-value | R^2^ | P-value |
| Elevation | 0.035 | 1.0 | 0.035 | 1.0 | 0.032 | 1.0 |
| Texture | 0.053 | 1.0 | 0.072 | 1.0 | 0.45 | **0.002** |
| Fine sand | 0.011 | 1.0 | 0.003 | 1.0 | 0.13 | 1.0 |
| Sand | 0.15 | 1.0 | 0.22 | 0.24 | 0.15 | 0.93 |
| Silt | 0.083 | 1.0 | 0.083 | 1.0 | 0.038 | 1.0 |
| Clay | 0.14 | 1.0 | 0.25 | 0.13 | 0.20 | 0.30 |
| Ammonium nitrogen | 0.068 | 1.0 | 0.035 | 1.0 | 0.022 | 1.0 |
| Nitrate nitrogen | 0.039 | 1.0 | 0.084 | 1.0 | 0.13 | 1.0 |
| Phosphorus | 0.16 | 0.89 | 0.30 | **0.024** | 0.30 | **0.026** |
| Potassium | 0.33 | **0.012** | 0.40 | **0.002** | 0.43 | **0.002** |
| Sulphur | 0.061 | 1.0 | 0.093 | 1.0 | 0.049 | 1.0 |
| Organic carbon | 0.034 | 1.0 | 0.072 | 1.0 | 0.027 | 1.0 |
| Conductivity | 0.16 | 0.87 | 0.19 | 0.39 | 0.22 | 0.19 |
| pH | 0.74 | **0.002** | 0.56 | **0.002** | 0.40 | **0.002** |
| Copper | 0.21 | 0.15 | 0.22 | 0.091 | 0.01 | 1.0 |
| Iron | 0.49 | **0.002** | 0.26 | 0.067 | 0.33 | **0.024** |
| Manganese | 0.095 | 1.0 | 0.16 | 0.86 | 0.14 | 1.0 |
| Zinc | 0.16 | 0.77 | 0.16 | 0.97 | 0.064 | 1.0 |
| Aluminium | 0.38 | **0.002** | 0.36 | **0.010** | 0.077 | 1.0 |
| Calcium | 0.53 | **0.002** | 0.54 | **0.002** | 0.32 | **0.019** |
| Magnesium | 0.14 | 1.0 | 0.19 | 0.51 | 0.12 | 1.0 |
| Sodium | 0.030 | 1.0 | 0.05 | 1.0 | 0.14 | 1.0 |
| Boron | 0.24 | 0.11 | 0.38 | **0.007** | 0.16 | 0.77 |
| Population density | 0.16 | 0.89 | 0.12 | 1.0 | 0.16 | 0.82 |
